# Combined docking and machine learning identifies key molecular determinants of ligand pharmacological activity on β2 adrenoceptor

**DOI:** 10.1101/2021.03.18.434755

**Authors:** Mireia Jiménez-Rosés, Bradley Angus Morgan, Maria Jimenez Sigstad, T.D. Zoe Tran, Rohini Srivastava, Asuman Bunsuz, Leire Borrega-Román, Pattarin Hompluem, Sean A. Cullum, Clare R. Harwood, Eline J. Koers, David A. Sykes, Iain B. Styles, Dmitry B. Veprintsev

## Abstract

G protein coupled receptors (GPCRs) are valuable therapeutic targets for many diseases. A central question of GPCR drug discovery is to understand what determines the agonism or antagonism of ligands which bind them. Ligands exert their action via the interactions in the ligand binding pocket. We hypothesised that there is a common set of receptor interactions made by ligands of diverse structures that mediate their action and that among a large dataset of different ligands, the functionally important interactions will be over-represented. We computationally docked ~2700 known β_2_AR ligands to multiple β_2_AR structures, generating ca 75,000 docking poses and predicted all atomic interactions between the receptor and the ligand. We used machine learning (ML) techniques to identify specific interactions that correlate with the agonist or antagonist activity of these ligands. The interpretation of ML analysis in human understandable form allowed us to construct an exquisitely detailed structure-activity relationship that identifies small changes to the ligands that invert their activity and thus helps to guide the drug discovery process. This approach can be readily applied to any drug target.

## Introduction

G-protein-coupled receptors (GPCRs) remain a therapeutically important family of proteins with over 100 receptors targeted by 500 drugs approved for clinical use^1^. The human β_2_-adrenoceptor (β_2_AR) responds to stimulation by the endogenous agonist ligands adrenaline and noradrenaline by inducing Gs-mediated cAMP signalling and is a valuable target for small molecule smooth muscle relaxants used to treat asthma and other pulmonary diseases^2,3^. Endogenous agonist activity can be readily inhibited by so-called antagonist drugs that prevent receptor activation by occupying the binding pocket without activation and blocking agonist access. A large number of ligands have been developed to target β-adrenoceptors (βAR) over the last 60 years since the pioneering discovery of beta-blockers by Sir James Black^3–7^.

All GPCRs share a common architecture of a bundle of seven transmembrane helices (TMs), with the ligand binding pocket accessible from the extracellular space and an intracellular effector binding site that becomes available following transition into an active receptor conformation^8^. One of the key features of GPCRs is that they are highly dynamic and adopt many distinct conformations that are important for engagement of signalling partners, e.g. activation of the Gs protein or arrestins^9^. It is generally thought that ligands control GPCR activity by preferentially stabilising active or inactive conformations^10^. With 35 reported structures with 13 diverse ligands in inactive and active states reported, β_2_AR is one of the best studied GPCRs from a structural perspective.

Structure-based drug design has become an integral part of the modern drug discovery process. Approaches to link ligand structure to its activity are generally based on the ligand chemical structure (similar chemical structure have similar activity paradigm) or by considering the interactions between the ligand and the receptor. Structural Interaction Fingerprints that describe the interactions of ligands with proteins^11–13^ have proven to be a very successful approach to score binding poses of ligands. A number of different interaction fingerprints have been developed, with more complex ones that incorporate atomic interactions and different types of non-covalent interactions having superior performance^14^. Several studies have attempted to link structural properties of the ligands and the interactions they make to the receptor to their functionality, based on available crystallographic structures and complemented with ligand docking^16,17^ or MD simulations^15^. These studies show significant promise in using interaction fingerprints to rationalise the link between structure and function, however the results of these studies were limited to the experimentally available structural data that cover only a very small fraction of known β_2_AR ligands. This limited their general ability to generate the new chemical knowledge needed to answer the key question in the drug discovery pipeline – what is the next molecule to make?

Ligands exert their action on GPCRs via the interactions they make in the ligand binding pocket. We hypothesised that despite the observed structural diversity of ligands targeting a particular receptor, there should be common interacting atoms within the ligand binding pocket that mediate their action. Given this hypothesis, we reasoned that among a large dataset of different ligands and their respective binding poses, the functionally important atomic interactions the ligands make with a particular receptor will be over-represented. To investigate this hypothesis, we assembled a database of ~2700 known β_2_AR ligands and computationally docked them to multiple experimentally determined β_2_AR structures, generating ca 75,000 docking poses (Figure 1A and B). For each of the docking poses, we generated a detailed Atomic Interaction Fingerprint (AIF), which comprises of a list of all the pairs of atoms involved in the interaction between a receptor and a ligand and a classification of each pairwise interaction as one of fifteen types of bond. In total, there were ca 1,100 possible interaction descriptors that we interchangeably call features (Figure 1C) in our dataset. Using pairwise correlation and Machine Learning (ML) approaches, we identified specific interactions between the ligands and the β_2_AR that correlated with their reported agonist or antagonist activity at the receptor (Table S1). In addition to a common set of interactions that were present for both ligand types, agonists make specific contacts with the amino acid residues H93^2.64×63^, K97^2.68×67^, S203^5.42×43^, S204^5.43×44^, S207^5.46×461^, H296^6.58×58^ and K305^7.35×34^ in transmembrane helices TM2, TM5, TM6, TM7 while antagonists make specific interactions with W286^6.48×48^ and Y316^7.43×42^ in TM6 and TM7. This approach successfully identifies the key features of the ligands in terms of the individual interactions they make with the receptor to exert their pharmacological action. Importantly, we were able to discover more subtle relationships where small changes to the ligand result in significant changes to their pharmacology, the so called *activity cliffs* encountered in every drug discovery program. This method represents a novel strategy for understanding the molecular mechanism of drug action on receptors and provides a valuable tool to guide the drug design process.

**Figure 1.**
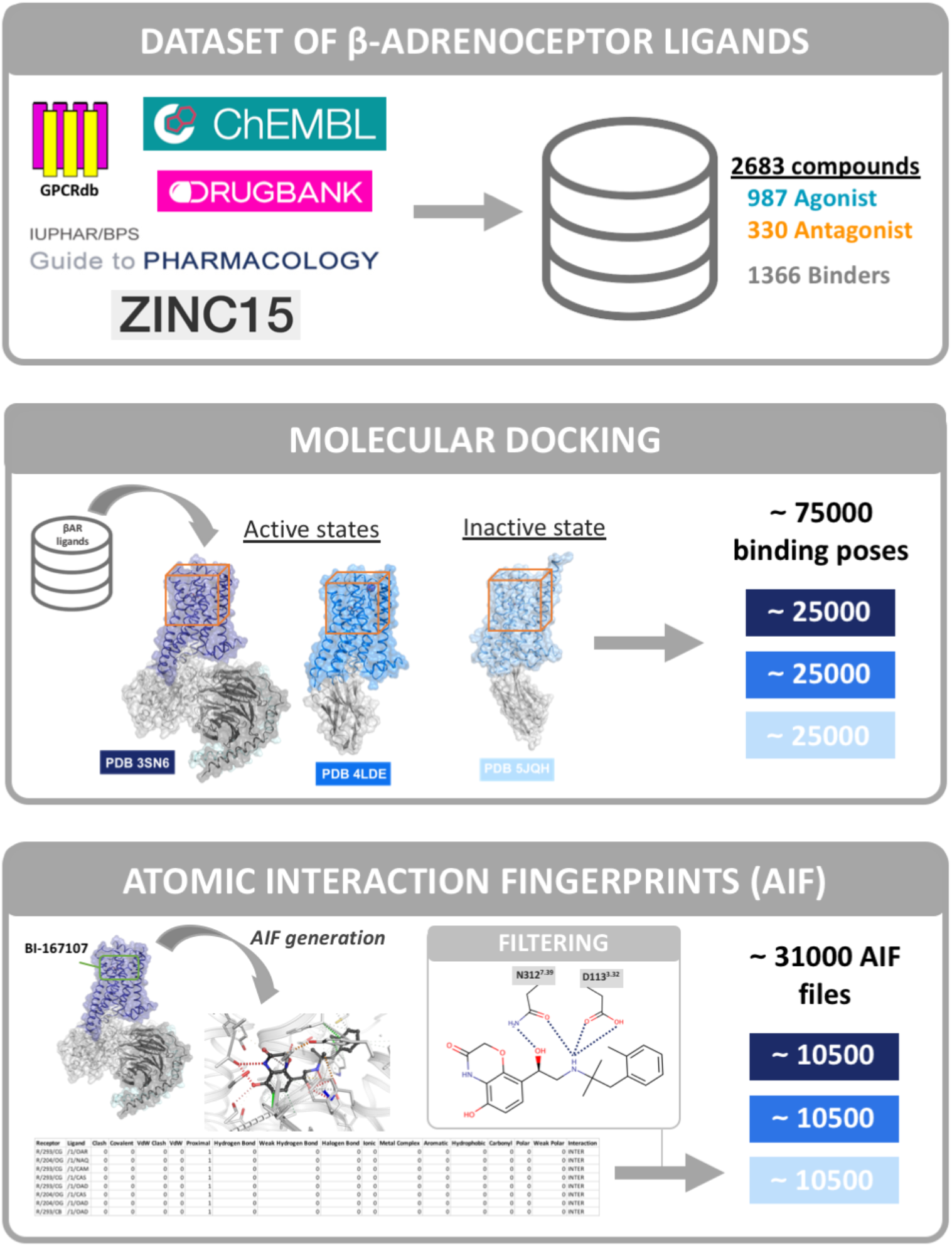
Workflow of the project. (A) Source of beta-adrenoceptor ligands available at open access repositories which comprise our 2683 compound dataset. (B) Molecular docking of test ligands to active and inactive β_2_AR structures was performed using Autodock Vina. (C) Interatomic interaction fingerprint (AIF) calculations were made using Arpeggio. In case of the “filtered dataset”, the generated AIFs were filtered based on the presence of ionic interactions with D113^3.32×32^ and N312^7.39×38^.

## Results

### β_2_AR agonists are on average bigger and more lipophilic compared to antagonists

To construct our dataset of currently known β_2_AR ligands we searched all available open access repositories such as GPCRdb, ChEMBL, DrugBank, Guide to Pharmacology and ZINC. The curated database included 2683 unique β_2_AR ligands, of which 1317 had reported pharmacological action (987 agonists and 330 antagonists/ inverse agonists). The remaining 1366 were classified as “known binders” with no assigned pharmacological activity (Figure 1A).

To understand if there are any obvious differences between agonists and antagonists, we compared their physicochemical (PC) properties predicted using OpenBabel software^18^. We found that many PC property values for agonists were statistically different from those for antagonists (unpaired t-test, p < 0.0001), for example, molecular weight (MW) and lipophilicity (logP) (Figure 2A and B, respectively). The MW of ~70% of agonist ligands was in the 350-550 g/mol range, with an average of 469 ± 108 g/mol. In contrast, the antagonist ligands were typically smaller, with ~70% within a range of 200-400 g/mol (average 358 ± 108 g/mol). The logP values of ~70% of agonists are in the range 3-7, with an average of 4.6 ± 1.6, whereas ~70% of antagonist ligands had logP values in the range 0-5 (average 3.1 ± 1.4). Taken together, the β_2_AR agonists profiled here tended to be more lipophilic and bigger in size. On the other hand, endogenous agonists adrenaline and noradrenaline are small and water soluble, suggesting that size and lipophilicity are not an intrinsic prerequisite of all agonists. We observed an identical linear correlation between the molecular weight and lipophilicity for both agonists and antagonists (Figure 2C), suggesting that that bigger compounds are more lipophilic. The likely explanation is that drug discovery efforts have focused on developing β_2_AR agonists formulated for the treatment of asthma. They are delivered to the lungs via inhalation with higher hydrophobicity increasing their duration of action at the target tissue. Therefore, although the observed differences in size and hydrophobicity are present in our data set, they are unlikely to have a functional role.

**Figure 2.**
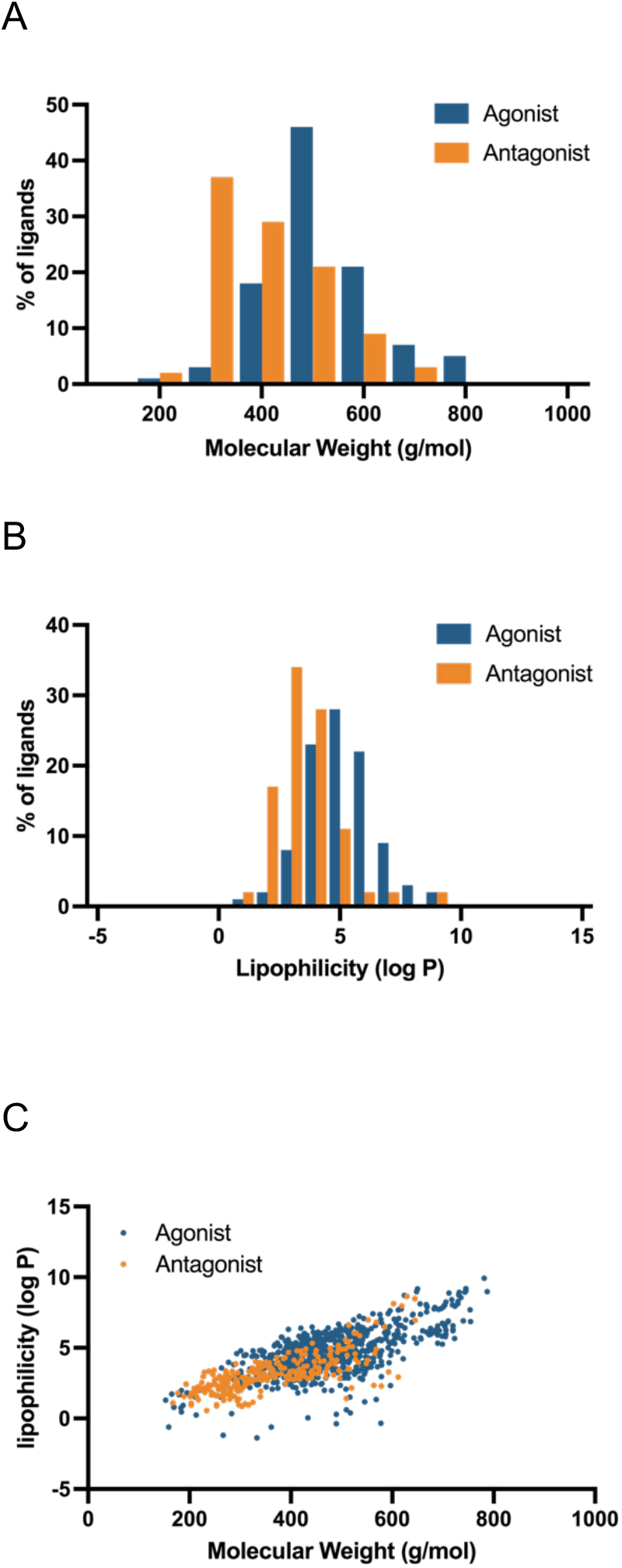
Physicochemical (PC) properties of the ligands (agonist in blue and antagonist in orange) predicted using OpenBabel software. (A) Molecular weight (MW) in g/mol, (B) lipophilicity (log P) and (C) correlation between lipophilicity and molecular weight. Spearman correlation coefficient is 0.62 for agonist and 0.76 for antagonists.

### Generating Atomic Interaction Fingerprints based on Molecular Docking Poses

To obtain structural information on how ligands in the curated dataset interact with the receptor (ie, ligand binding poses), we performed molecular docking using the open-source AutoDock Vina software^19^. Three β_2_AR structures were studied: the active conformational states i) PDB 3SN6 stabilised by the Gs protein^20^ and ii) PDB 4LDE stabilised by a nanobody^21^, and iii) the inactive conformational state PDB 5JQH^22^. We obtained ~75,000 binding poses in total, ~25,000 poses for each PDB (up to 10 poses for ligand, for 2683 compounds) (Figure 1B). Each ligand binding pose was used to generate an atomic interaction fingerprint (AIF) using Arpeggio software^23^, in total we obtained ~75,000 AIF files (Figure 1C). Each AIF included ~60 unique interactions on average between the atoms of the ligand and atoms of the receptor. When the type of atoms of each ligand and the type of bond formed are considered, this resulted in over 1,100 possible types of interaction across the complete ligand dataset.

It is important to consider that the obtained AIF fingerprint dataset contains noise because not all of the predicted docking poses are likely to be relevant or functionally important. The limitations of the ligand docking algorithms result in multiple alternative binding poses with very similar “quality scores”, with only one of the top ten solutions likely to correspond to the experimentally observed binding pose. While crystallographic structures typically represent one ligand binding pose, they tend to represent the lowest energy state of the system. On the other hand, molecular dynamics simulation and biophysical experiments suggest that ligands are dynamic when bound to the receptor^24^. Therefore, it is important to consider multiple ligand docking poses in the analysis. We rationalised that in a large dataset of different ligands and their respective binding poses, the functionally important atomic interactions between the ligands and the receptor will be over-represented while the influence of the noise (irrelevant binding poses) would average out.

We improved the signal-to-noise ratio within our dataset by excluding irrelevant binding poses using prior knowledge based on crystallographic data (Figure 1C, filtering panel). The majority (ca 97%) of β_2_AR ligands have a prevalent β-hydroxy-amine motif that makes specific interactions with the receptor. We therefore excluded poses that did not display this ionic interaction between the oxygen of D113^3.32×32^ and the nitrogen atom of ethanolamine of the ligands and the hydrogen bond between the oxygen atom of N312^7.39×38^ and either the NH or beta-hydroxyl groups in the ligand scaffold; these have been observed in every experimental crystallographic structure of the β_2_AR. After applying this filter, we obtained ~31,500 atomic interactions files (~10,500 poses and AIF files for each PDB), reducing the size of the original dataset by ~55%. We refer to this as the “filtered dataset”. As the filtering step also removed ~3% of ligands in our dataset that did not contain the β-hydroxy-amine motif or did not produce suitable poses, we have also included in our analysis the “full dataset” consisting of ~75,000 AIF files with no filtering for comparison.

### Data-driven analysis reveals key interactions that drive agonism and antagonism of ligands

We constructed a ligand-receptor interaction matrix, organising the atom-atom interactions and their types in the columns and each binding pose in rows for each PDB. We defined the ligand binding site as all residues that interact with at least one ligand binding pose in the dataset resulting in 30 residues in total (Table S2). The atoms of the ligand binding site provide a constant reference coordinate system to describe ligand-receptor interactions. We defined atomic interaction between specific atoms of the receptor, and the specific atom (C, N, O, etc) in the ligand and the nature of the interacting bond (polar, ionic, hydrophobic, etc). This strategy allowed us to encode the ligand-receptor interaction matrix that accommodates diverse ligands irrespective of their structural scaffold.

Using Pearson’s pairwise correlation between the independent variables describing the presence or absence of an atomic interaction and the dependent variable denoting agonist/antagonist properties of the ligands, we identified atom-atom interactions (or features) that associated with agonism or antagonism in the filtered dataset. From about 100 commonly observed interactions, we find that the most representative interactions for agonist ligands are hydrophobic/aromatic contacts involving K97^2.68×67^, F194^ECL2^, H296^6.58×58^ and K305^7.35×34^ and polar/ionic/hydrogen bond contacts with S203^5.42×43^, S204^5.43×44^, S207^5.46×641^ and H296^6.58×58^. The antagonists made specific hydrophobic/aromatic contacts with W286^6.48×48^ and Y316^7.43×42^ and polar/ionic/hydrogen bond contacts with Y316^7.43×42^ (Figure 3A and Table S3).

**Figure 3.**
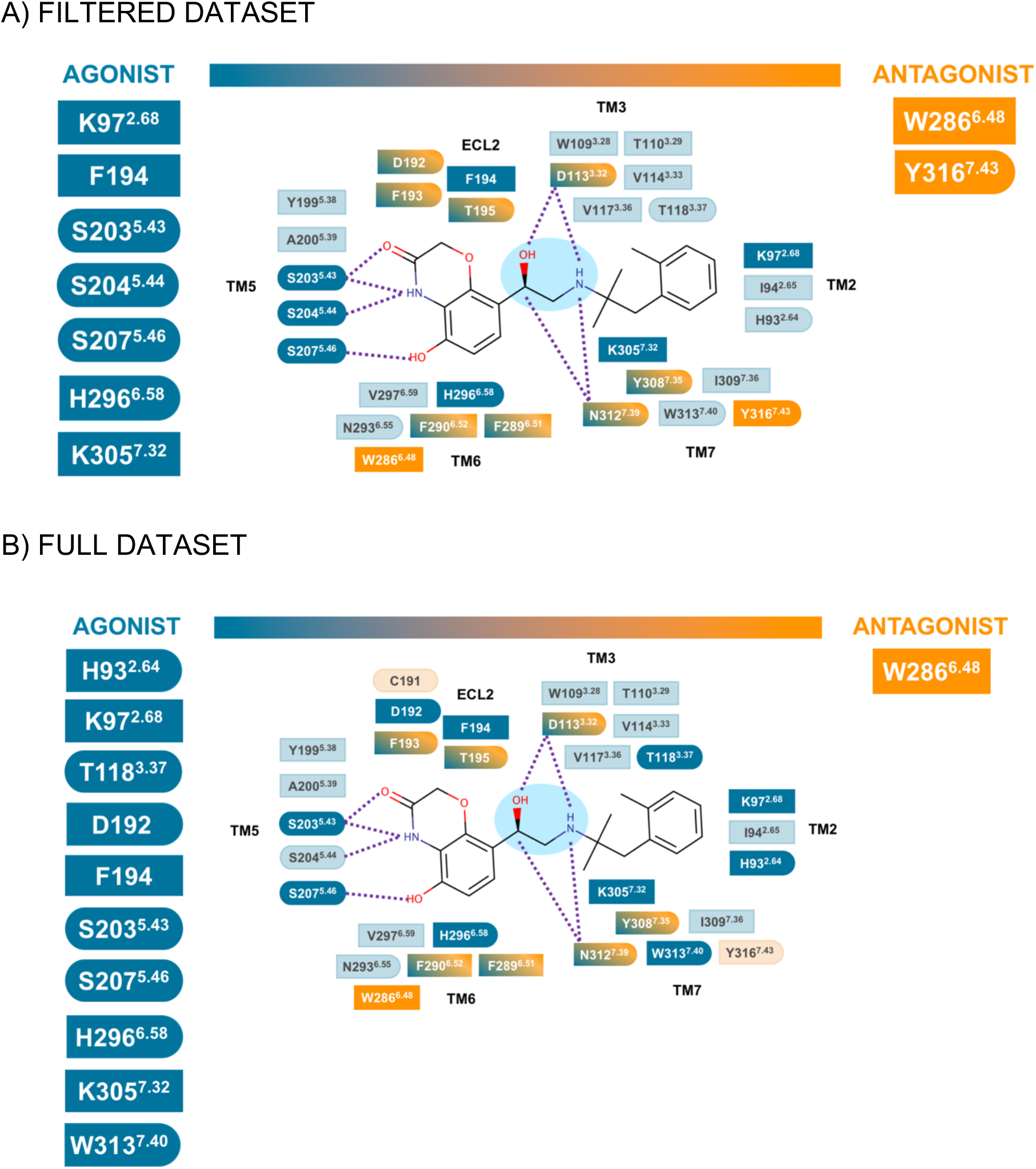
Schematic representation of the interactions predicted using the pairwise correlation approach. (A) filtered dataset and (B) full dataset. The type of interaction is summarised in squared shape for hydrophobic and aromatic contacts, round shape for the polar, ionic and hydrogen bond contacts, and the combination of both. The dotted purple lines represent ionic and/or hydrogen bond contacts. The ethanolamine moiety of the BI-167107 ligands is highlighted in light blue.

While the majority of interactions had the same impact on receptor function (mediating agonism or antagonism) for all atoms of the individual residue, in some cases (D113^3.32×32^, D192^45.51×51^, F193^45.52×52^, T195 ^ECL2^, F289^6.51×51^, F290^6.52×52^, Y308^7.35×34^, N312^7.39×38^) this depended on the individual atoms of the residue and the nature of the interacting bond (Table S3). For example, the polar/ionic/hydrogen contact of the carbonyl oxygen (OD1, as defined by the Protein Data Bank format^25^) of D113^3.32×32^ with an oxygen atom of a ligand is predictive of agonism while interaction with a nitrogen atom is predictive of antagonism. Contacts made by hydroxyl oxygen (OD2) of D113^3.32×32^ have the opposite effect: interaction with a nitrogen atom of the ligand corresponds to agonism, whilst interaction with an oxygen atom results in antagonism. In another example, polar contacts of the sidechain nitrogen (ND2) of N312^7.39×38^ with oxygen atoms in the ligand corresponded to agonism whilst interaction with nitrogen leads to antagonism.

The full dataset was a more complex challenge as it contains more noise in terms of the number of different poses and also a more diverse range of ligands. Nonetheless we also observed around 100 common interactions, which were mostly the same as those determined for the filtered dataset. However, several interactions changed their relative importance (Figure 3B); for example, the importance of S204^5.43×44^ as a determinant of agonism was reduced, while W313^7.40×39^ became more predictive of agonism. However, the core set of agonist-associated interactions made with S203^5.42×43^, S207^5.46×461^ and, F194^ECL2^, H296^6.58×58^, K305^7.32×31^ and K97^2.68×67^ remained the same.

To validate the performance of the Pearson’s pairwise correlation, we computed the maximum Matthews Correlation Coefficient (MCC) which measures the quality of binary classifications when the classes are of different sizes as in our case (ca 75% are agonists). For the filtered dataset, taking the maximum MCC with a cut-off score of 0.37, we obtained a pharmacological classification (agonist or antagonist) with a MCC of 0.43 that corresponds to the accuracy of prediction of 79% (Figure S1A). For the full dataset (cut-off = 0.51), the MCC and accuracy decreased to 0.29 and 67%, respectively (Figure S1B). An important consideration for interpretation of the prediction accuracy is that the training dataset may contain errors: compounds that are “wrongly” assigned to a particular class (e.g., agonist or antagonist). Therefore, we would not expect the predictors to be 100% accurate during the validation step.

As the pairwise correlation approach identifies the relative importance of individual interactions, we applied ML strategies (see methods for details) that can detect more complex patterns in the data than pairwise correlation analysis. We trained a Random Forest Classifier (RFC)^26^ on the filtered dataset and XGBoost^27^ on the full dataset. RFC constructs a multitude of decision trees and averages them to improve the predictive performance and control overfitting, reaching MCC values in training of 0.81 and an accuracy of 92% on the filtered dataset (Figure S2 and S4A). The XGBoost algorithm that iteratively constructs optimised decision trees guided by the results of the previous steps performed remarkably well on the full dataset (Figure S3 and S4B), with a prediction performance on the holdout set of 0.78 MCC and 93% accuracy after full Bayesian optimisation. This suggests that there are predictive patterns in both the filtered and full dataset not captured by a simple predictor based on pairwise correlations.

It is, however, a considerable challenge to interpret what the ML algorithms have actually learned. We extracted the feature importance for RFC trained on the filtered dataset (Figure 4A,B) and the feature importance for XGBoost trained on the full dataset (Figure 4C,D), using the Shapley Additive Explanations (SHAP) values which reflect the contribution of each feature to the prediction. In most cases, the presence of a particular interaction is predictive of agonism or antagonism. However, in a minority of cases, the absence of the interaction was more important for predictions (e.g., 193/CB-1/C hydrophobic).

**Figure 4.**
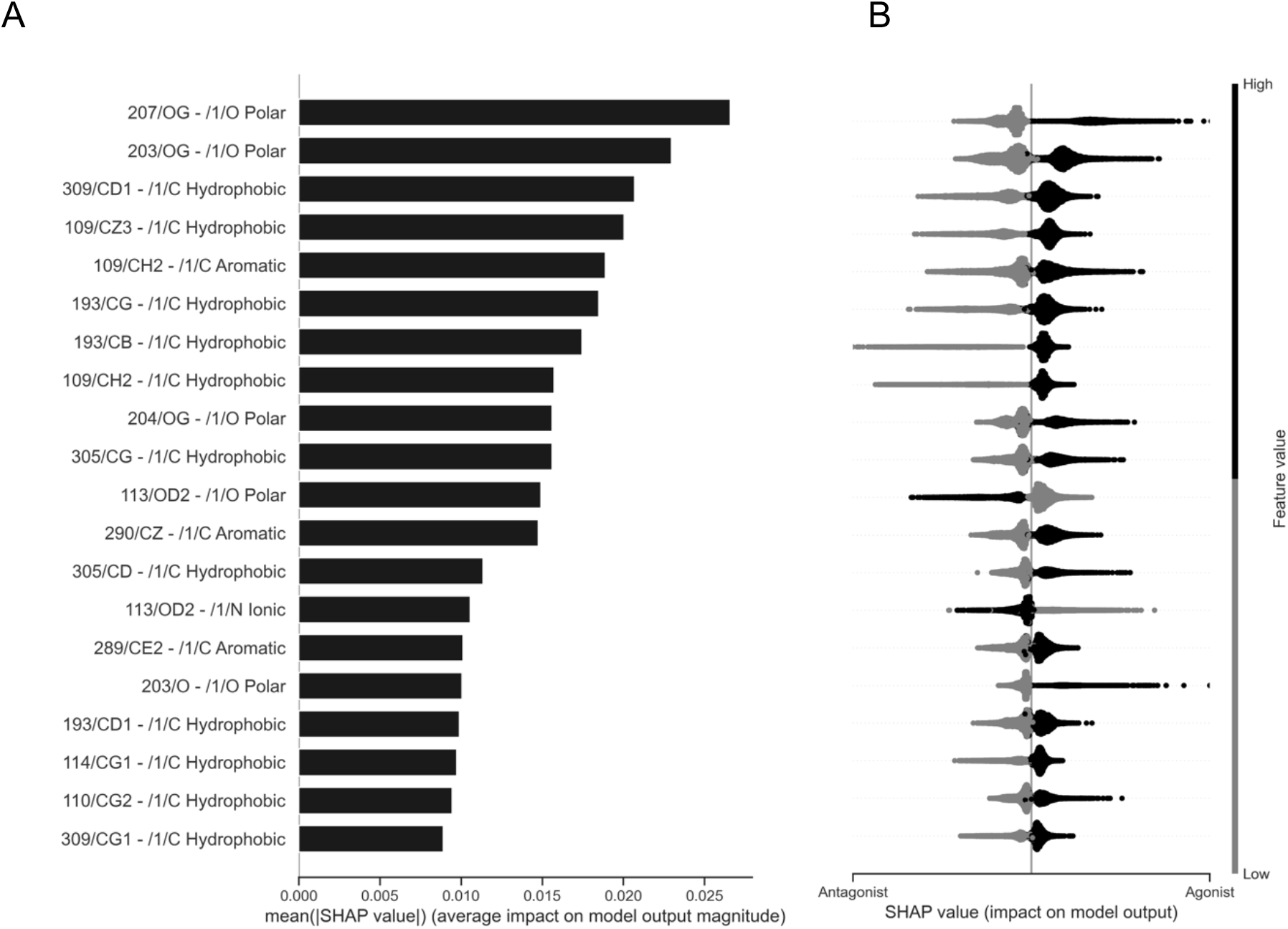
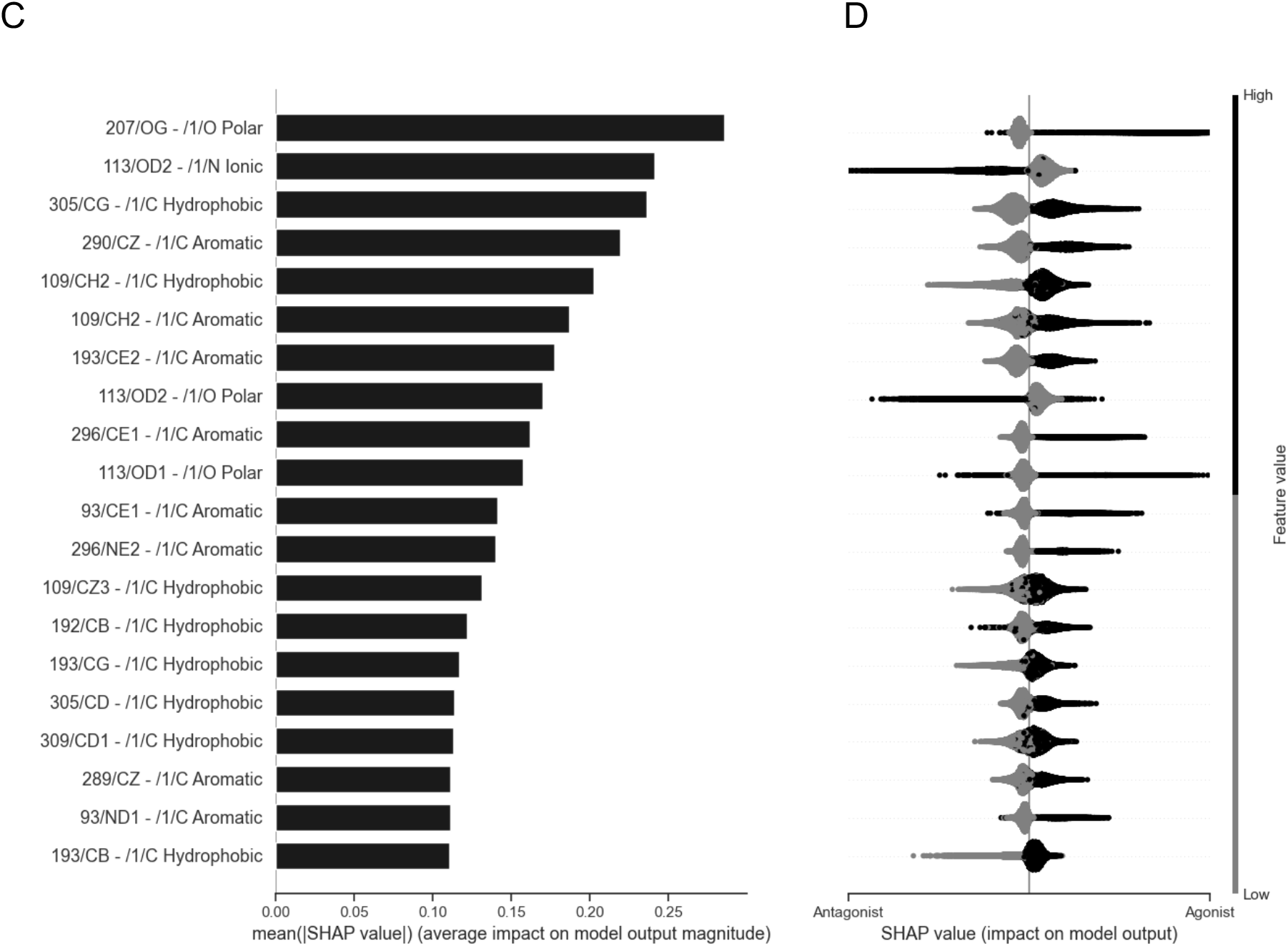
Feature importance of the RFC (A,B) and XGBoost ML (C,D) models applying SHAP value method. (A,C) The x-axis is the average magnitude change in model output when a feature is “hidden” from the model. Higher SHAP values indicate higher importance of the feature. (B,D) Local SHAP values per sample (each ligand pose) sorted by the mean absolute SHAP value method. Grey represents a value of 0, thus indicating the absence of a particular atomic interaction for a specific sample. Black represents a value of 1, thus indicating the presence of a particular atomic interaction for a specific sample. The x-axis shows how the presence or absence of an atomic feature increases or decreases the likelihood of a sample being classified as an antagonist. The data are plotted for all samples in the dataset, showing the distribution of importance values. The units of the x-axis using RFC and XGBoost are log odds.

Overall, while the relative order of importance of individual features varied depending on the model, we observed the same set of interactions that were predictive of agonism or antagonism for both models (Table S4). The application of pairwise correlation analysis and ML methods allowed us to identify the key interactions associated with agonism or antagonism of ligands (Figures 3 and 5).

**Figure 5.**
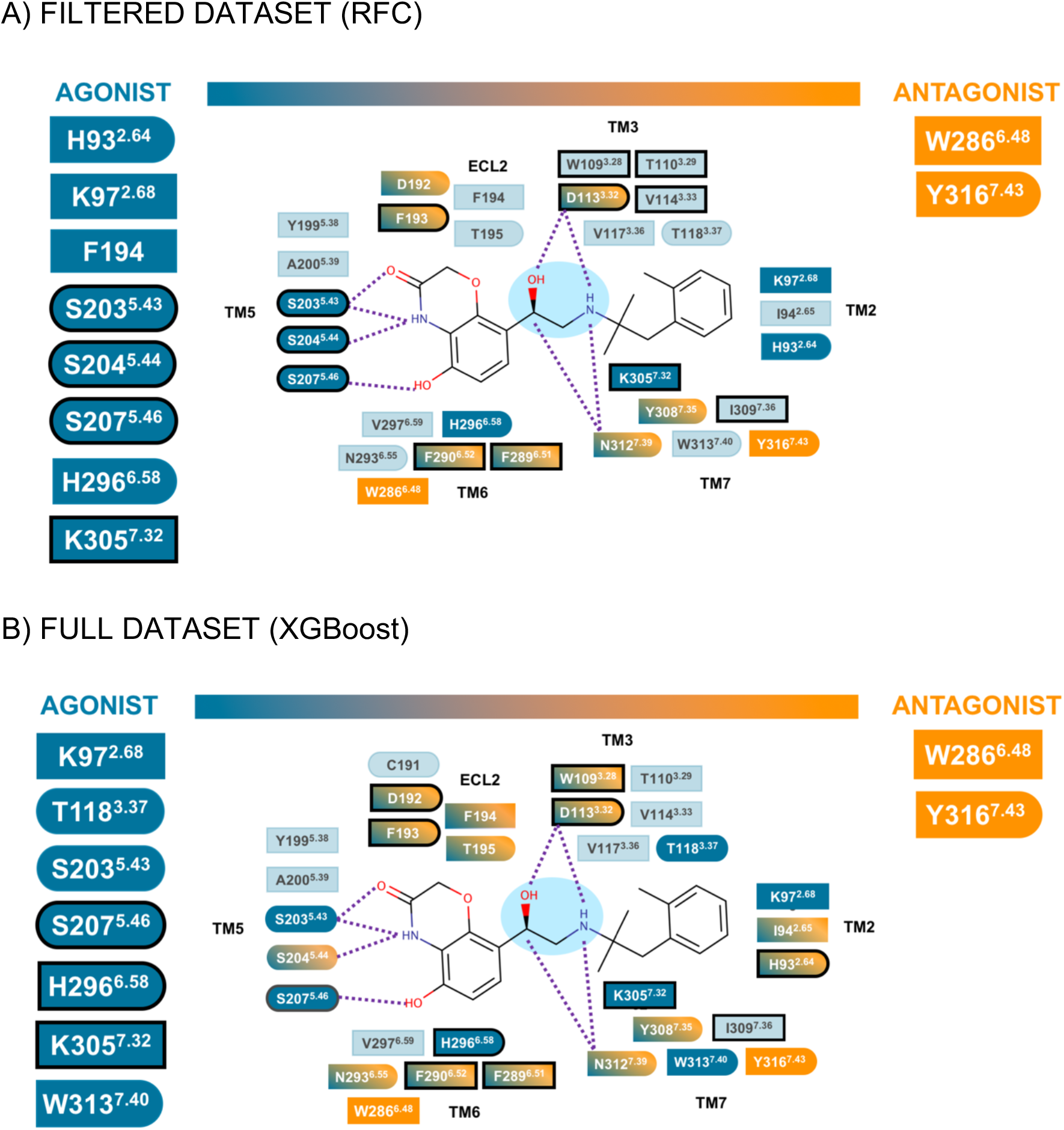
Schematic representation of the interactions for the machine learning approach. (A) RFC for the filtered dataset and (B) XGBoost for full dataset. The type of interaction is summarised in squared shape for hydrophobic and aromatic contacts, round shape for the polar, ionic and hydrogen bond contacts, and the combination of both. The dotted purple lines represent ionic and/or hydrogen bond contacts. The black outlines represent the atomic interactions with higher feature importance. The ethanolamine moiety of the BI-167107 ligands is highlighted in light blue.

## Discussion

While an observation that on average agonists are larger and more hydrophobic could potentially be used to distinguish them from antagonists in βAR ligand dataset, the pharmacological action of ligands on GPCRs is far more specific than a simple function of their size or hydrophobicity.

### Specific ligand-receptor interactions determine their pharmacological activity

While ML algorithms can successfully classify compounds into agonists and antagonists, understanding what their decision is based on and translating this information into the language humans can understand is crucial for their usefulness for drug discovery^28^. Studying the ligand binding poses of thousands of ligands docked in the β_2_AR binding pocket allowed us to identify the key ligand-receptor interactions which dictate a molecule’s propensity to cause agonism or antagonism. The structurally diverse nature of the test set that consisted of all ligands with reported activity in publicly accessible databases allowed us to identify several “hot spots” mediating the agonism or antagonism of ligands acting on β_2_AR. Agonism was mediated by residues in TM2 and TM5, and further facilitated by residues in TM6 and TM7. It is entirely plausible that certain ligands can successfully pull these TM regions together causing receptor activation in the process. In contrast, our data suggest that antagonism is mediated by the interaction of ligands with W286^6.48×48^, the so called toggle switch, that has long been proposed to play a key role in the activation of GPCRs^29,30^. The second mediator of antagonism is Y316^7.43×42^ which is involved in the so called 3-7 lock that has previously been identified as important for GPCR activation^31^. Engaging these key residues in the ligand binding pocket likely prevent the conformational rearrangements necessary for activation of the receptor.

### Potential for developing more fine-grained models of ligand activity

While the assembled data classify compounds as agonist or antagonist, the pharmacological activity of compounds covers a spectrum from a very strong antagonist (aka inverse agonist) to that of a very strong agonist (aka full agonist). Another class of GPCR ligands, so called biased ligands, changes the balance between activating G protein and arrestin signalling pathways, with a potential to increase their therapeutic benefits^9,32^. It is likely that such partial and biased ligands would also show a distinct AIF that is somewhat different from the all-inclusive agonist AIF we have identified in the current work. However, a large experimental dataset of partial or biased agonists would be needed to explore this hypothesis, ideally collected in a uniform screen to minimise experimental and interpretational bias. The analysis of the learning performance of RFC and XGBoost classifiers (Figure S4) suggest that reasonable performance is achieved with a limited dataset (ca 300-450 compounds), although further increases in the dataset size resulted in improved performance. It is likely that an even larger dataset would be required to predict continuous rather than binary structure-activity relationship from AIFs.

Our methodology can be readily applied to any receptor (or drug target) for which an extensive set of ligands has been developed and characterised, and where *in silico* docking experiments can be performed. This can include data already in the public domain or through examining the results of an *in house* (*e.g., commercial*) drug-target screening campaign. The advantage here is that in many cases the same signalling assay will have been used to profile all the compounds, improving the consistency of the dataset. This would allow the relative importance of each atom-atom interaction to be assessed as a modifier of signalling output. Also, it may be possible to isolate functional readouts (e.g. β–arrestin versus G protein) and therefore make predictions about functional bias. Further tantalising possibilities include the use of automated internet meta search of publications and patents to assemble such datasets and reduce the number of compounds described as “known binders” if they are not available yet.

### Potential for developing predictors of pharmacological activity for novel ligands

Being able to understand which atoms of the ligand drive agonist or antagonist activity significantly increases the value of *in silico* docking campaigns. Importantly, it opens doors to a more rational engineering of ligands with improved and optimised pharmacological properties – facilitating the design of new ligands not present in the large virtual libraries and thus opening up a chemical space many orders of magnitude larger than the largest virtual libraries available.

From a computation perspective, it is a relatively straight-forward task to generate a prediction of ligand pharmacological activity based on the model learned and the predicted binding pose of the ligand and the corresponding AIF. However, large scale docking experiments produce multiple possible ligand binding poses, and the existing scoring functions do not allow for reliable identification of the “correct” binding pose. The structural diversity of the ligands complicates the analysis even further as overlaying of the predicted binding pose with the available experimental data is not always informative.

Our data strongly support the hypothesis that individual atomic interactions are correlated with ligand pharmacological activity. This is learned from a large dataset of ligand binding poses, where “correct” binding poses are a minority but machine learning methods we used identified the structure-activity relationship because “wrong” binding poses averaged themselves out. Prediction of pharmacological activity, on the other hand, is 100% dependent on having a correct binding pose for the ligand. This is a problem that has not yet been solved in a satisfactory manner, and it limits the performance of any structure-based activity prediction method. It is clear that the future progress in our ability to predict the pharmacological activity of novel ligands will be closely correlated with our ability to correctly predict their ligand binding poses.

## Conclusions

These results strongly support the hypothesis that the interatomic interactions between the receptor and its ligands are central to differentiate between their agonist and antagonist effects at the β_2_AR. The overview obtained of the interatomic interactions between receptor and ligand which correlates with an action will help the synthesis of new previously unseen compounds with a specific pharmacological activity. The growth of GPCR ligand databases provides a rich data source to facilitate the application of this approach to other GPCRs, while conceptually this approach could be applied to any drug target.

The ability to predict the pharmacological action of a ligand based on its ligand binding pose will significantly advance drug discovery projects contributing to a reduction of attrition during drug development. The tools presented have the potential to focus the efforts of chemists proposing new candidate molecules based on existing scaffolds and offers the opportunity to identify completely new scaffolds that may be more amendable to modifications from large scale docking experiments, thus opening up a chemical space many orders of magnitude larger than the largest virtual libraries available.

## Acknowledgements

We thank Michael Stokes, Jill Baker, Uwe Grether, Arne Rufer and Wolfgang Guba for valuable advice and stimulating discussions. MJR was funded by COMPARE. BAM, CRH and SAC are funded by the MRC IMPACT doctoral programme. DAS is funded by the Roche Postdoctoral Fellowship. PH is funded by an international fellowship awarded by the Office of Educational Affairs, Thailand. AB is funded by the Study Abroad Postgraduate Education Scholarship (YLSY) awarded by the Republic of Turkey Ministry of National Education. LBR was funded by EMBO short term fellowship and the COST action 18133 ERNEST short term scientific mission fellowship. MJS was funded by the BBSRC Doctoral Training Programme, University of Nottingham and COMPARE.

## Methods

### Dataset preparation

A dataset was compiled using the primary open access repositories GPCRdb^1,2^, ChEMBL^3^, ZINC^4^, DrugBank^5^ and Guide to Pharmacology^6^. This dataset yielded a total of 2,643 unique β_2_AR ligands, of which 1317 have reported pharmacological action, while 1,326 compounds are binders with undetermined activity profiles. We classify ligands with known activity as either agonists (including partial and full agonists) or antagonists (including inverse agonists).

Each ligand was assigned an internal ID (ranging from 1 to 2,643), and its corresponding SMILES string (line notation encoding its molecular structure) and pharmacological action (agonist/antagonist/binder) were retrieved from the relevant databases. The International Chemical Identifier key (InChIKey) was used as a unique identifier to distinguish between ligands across the dataset^7^. Both InChIKey and physicochemical properties appended for all compounds were acquired using the software Open Babel v3.1.1^8^.

### Protein structures and ligand preparation and Docking

The active-state protein coordinates were extracted from two crystal structures of human β_2_AR bound to an ultrahigh-affinity agonist (BI167107) coupled with the Gs protein^9^ or/and with a G protein-mimicking nanobody (Nb6B9)^10^ from the Protein Data Bank (PDB code: 3SN6 and 4LDE, respectively). The inactive-state protein coordinates were extracted from the human β_2_AR bound to the inverse agonist carazolol (PDB code: 5JQH)^11^.

Receptor structures were aligned to use the same grid box of 22 × 22 × 32 Å at the orthosteric binding site, protonated and charged, yielding a protein input file for subsequent docking experiments using UCSF Chimera^12^.

The SMILES representation of ligands along with their internal ID were protonated and converted to a spatial data file (SDF) and pdbqt formats using Obabel.

The semi-flexible molecular docking was carried out using the software AutoDock Vina^13^ and generated up to 10 poses for every compound. In total, 2,643 compounds were docked in three β-adrenoceptor structures, yielding almost 27,000 docking poses.

### Interaction fingerprint calculations and filtering

The inter-atomic receptor-ligand interaction fingerprints (AIFs) were calculated for all docking poses generated for each compound using the software Arpeggio^14^ executed in Docker environment^15^, a software container platform. This method accounts for the presence of up to 15 subtypes of interatomic interactions, classified by atom type, distance and angle constraints. The output was presented as binary values, with a 1 denoting the presence of a particular defined interaction and 0 indicating an absence. A Python script was written to filter the Arpeggio results (to generate the “filtered dataset”) by imposing minimum constraints that enforced certain features deemed essential for β_2_AR ligand binding, which it eliminated all irrelevant binding poses (around 50% rows). Criteria important for binding were based on prior knowledge derived from the literature, in particular the presence of the ionic/polar interaction between D113^3.32^ and N312^7.39^ with the ethanolamine moiety of the ligands.

### Generation of interaction matrix

A Python script was written to process each docking pose to generate a single *MxN* matrix for each PDB, where *M* are ligand poses (samples as ‘ligand internal ID_docking pose number’) and *N* are the specific atomic interaction (features as ‘receptor residue number/interacting atom – ligand interacting atom and interaction type’ (e.g. ‘lig 752_04’ and ‘301/O - N Polar’ respectively)), present in the whole ligand set. The value 1 corresponds to the occurrence of a particular type of interaction and 0 to the non-occurrence of a particular type of interaction. As the AIF files generated by Arpeggio only contained the interaction present for a particular docking pose, the imputation of missing data was handled by setting any undefined (NaN) values to 0. Finally, we excluded the 3 subtypes of interactions reported by Arpeggio: Clash, VdW Clash and Proximal from subsequent analysis as they provided very little information but represented around 60% of the columns. We also included pharmacological action label (“agonist” or “antagonist”, if known, or “binder” if not known) for each ligand binding pose in the same data table as an additional column.

### Descriptive statistical analysis

A descriptive statistical analysis of the frequency of the interatomic interactions was performed using a Python script. In this manner, the most frequently occurring features (those observed in at least 10% of all docking poses) contributing to agonism and antagonism were clearly identified across all interaction types and collated for further analysis. This resulted in the reduction of the number of features from ca 1,100 to ca 100.

Subsequently, we computed pairwise correlation between the columns representing atomic interactions and the column representing pharmacological action using Pearson’s correlation coefficient (r) method. The resulting value r for each interaction (feature) reflects how well it is correlated with the pharmacological action (agonism or antagonism). Plots, graphs, and tables were generated with Excel, and statistical significance was determined using an unpaired t-test using Prism 8.

### Machine learning dataset preparation

The dataset was then randomly shuffled and split, via stratification, into crossvalidation and final hold-out datasets. The cross-validation set was used for training and validation during hyperparameter optimisation. The hold-out dataset comprised 2% of the original dataset and allowed us to gauge whether the validation scores were good estimations of model performance when generalising to unseen data. The holdout set was not used during any training or optimisation procedures.

### Model Selection

#### Performance Metric

For the filtered dataset the Random Forest classifier and for the unfiltered dataset XGBoost classifier were used. Matthews correlation coefficient (MCC) was used as the performance metric for all models^16^. The MCC metric is defined as follows:

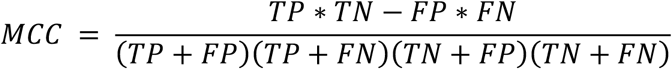

Where *TP* is the number of true positives, *TN* the number of true negatives, *FP* the number of false positives and FN the number of false negatives. The MCC for binary classification weights both positive and negative classes equally, whilst also being robust to severe class imbalances. A value of +1 indicates a perfect positive correlation, that is a total agreement between prediction and observation. An MCC score of 0 indicates no correlation, that is the classifier performs no better than a random coin flip. Finally, −1 indicates a perfect negative correlation, that is a total disagreement between prediction and observation.

#### Model Performance Estimation

Model performance was validated using repeated-stratified-k-fold cross-validation. Cross validation entails splitting the dataset into k equally sized partitions, termed folds. One of the folds is extracted and used for validating a model on unseen data. The remaining folds are then used to train the model. This process is then repeated using each of the k folds as the validation set. The optimal model is the one which has the best performance on average across all k folds. Cross validation generally provides a less optimistic estimation of model generalizability on unseen data, which is finally tested on the holdout set. Due to class imbalances in the data, stratification is used to ensure that the original distribution of classes is maintained in each fold, thus preventing any fold from being populated by a single class^17^. Model estimation can be noisy and so by performing cross-validation over many repeats one obtains a more precise estimation of true model performance. Bootstrap resampling was used to estimate model uncertainty^18^. Confidence intervals were calculated with respect to a 99% confidence level. Bayesian hyperparameter optimisation (BHO) was utilised to determine high performing model parameter configurations when tested on unseen data. BHO was set to maximize the mean MCC across K-folds and repeats, model uncertainty was then calculated using optimised models only.

##### Random Forest Classifier Hyper Parameters

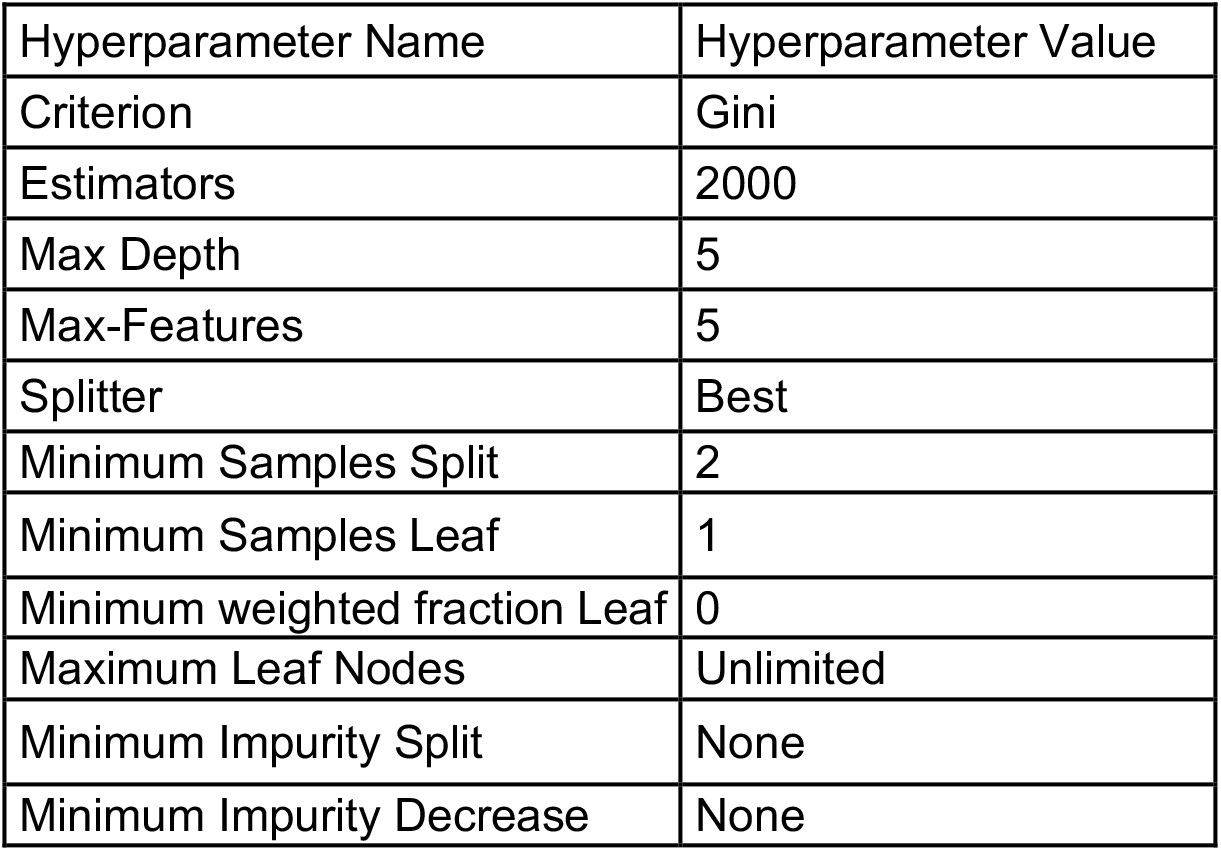

##### XGBoost Hyper Parameters

| Hyperparameter Name | Hyperparameter Value |
| --- | --- |
| Eta | 0.4306 |
| gamma | 0.2458 |
| Learning Rate | 0.05873 |
| Max Delta Step | 7 |
| Max Depth | 8 |
| Minimum Child Weight | 1.246 |
| Number of Estimators | 1150 |
| Scale Positive Weight | 1 |
| Subsample | 0.7532 |

All other hyperparameters for XGBoost that are not specified were kept at their default values according to the XGBoost API guide (https://xgboost.readthedocs.io/en/latest/parameter.html)

#### Model Feature Importance analysis

The most important atomic interactions, for classifying agonist or antagonist ligands, were identified using the Shapley Additive Explanations (SHAP) method^19^. Shapley values are based upon coalition game theory and inform one how to fairly distribute the prediction of a model among the features. The Shapley value for one feature is the average marginal contribution of a feature value across all the possible combinations of features. More concretely the Shapley value assigns an importance to each feature by calculating the effect on model prediction when including a particular feature compared to the model prediction when the feature is withheld. Mathematically this can be formalised as

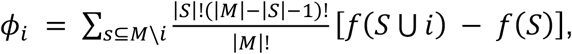

where *S* refers to a subset of features that does not contain the feature for which we are calculating *ϕ_i_. S* ∪ *i* is the subset that contains features in *S* and feature *i*. Finally, *S* ⊆ *M*/*i* represents all sets *S* that are subsets of the total set of features *M*, excluding feature *i*. The computation time increases exponentially with the number of features; thus we used the TreeSHAP algorithm that approximates SHAP values for tree-based machine learning models in polynomial time^20^. The main motivations for using the SHAP feature importance method over other popular methods, such as Gini and Permutation methods, is due to the following:

##### Consistency

The Gini feature importance method is susceptible to producing inconsistent feature importances that are biased to the specific ordering of features specified by their position, as split nodes, in the tree. TreeSHAP method is equivalent to averaging differences in model predictions over all possible orderings of the features and thus does not suffer from such inconsistencies.

##### Granular Interpretability

Although permutation importance is not biased to the specific structure of decision trees it only provides a global understanding of the most important features. With TreeSHAP, observations get their own set of SHAP values and therefore we can understand feature importance on a per sample basis.

#### Determining The Optimal Number of Repeats

There is an exponential relationship between the number of times one has to repeat Bootstrap or Cross-Validation and the level of precision to within which one would like to measure true model performance. This leads to a trade-off between the precision and time complexity of model performance estimation. We thus estimate the optimal number of repeats to use for Bootstrap and cross-validation resampling methods to an acceptable level of precision as:

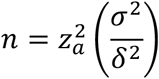

where *z* is the ordinate on the Normal distribution curve that corresponds to a particular level of confidence we have in our estimation, denoted *α. σ* is the population standard deviation and *δ* is the specified precision of the estimate. We estimate the population standard deviation via repeated bootstrap resampling, thus each estimate of the number of repeats is specific to the variance of each model and its hyperparameter configuration (https://www.itl.nist.gov/div898/handbook/ppc/section3/ppc333.htm).

A precision of 1% (Marginal Error = 0.01) was selected for all resampling methods (Supplementary Figure 5). Therefore, a minimum of 13 repeats for both the RFC and XGBoost were used during cross-validation and bootstrap resampling.

