## Supplementary material for "Combined docking and machine learning identifies key molecular determinants of ligand pharmacological activity on β2 adrenoceptor": SI

### Supplementary data

Supplementary Figure 1

B. FILTERED DATASET

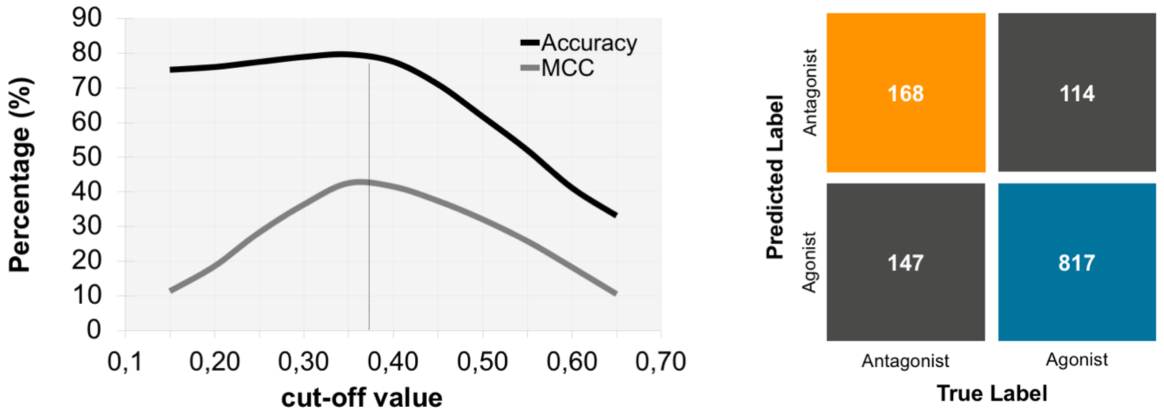


B. FULL DATASET

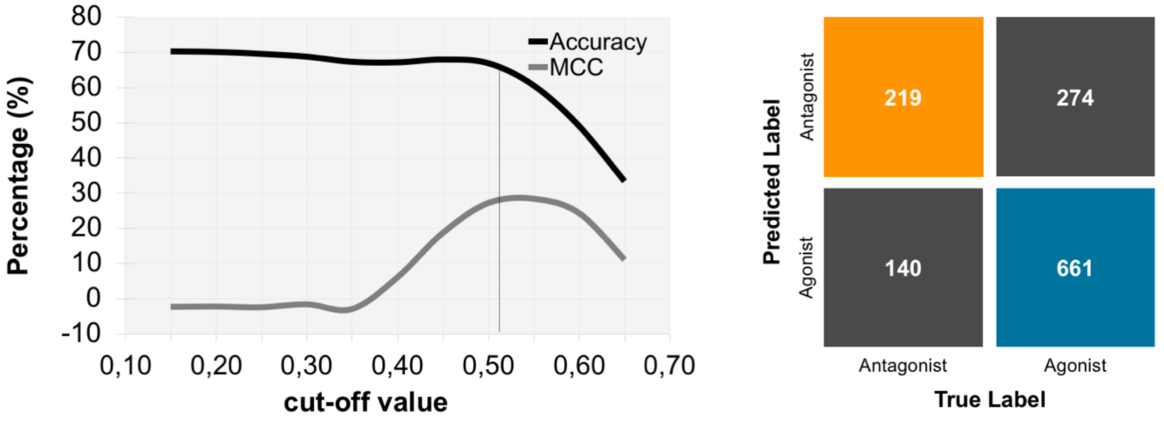


Supplementary Figure 1. Cut-off graph to obtain the maximum MCC value for the Pearson's correlation approach and the corresponding confusion matrix to differentiate agonist and antagonist. (A) Filtered dataset, right: cut-off and left: confusion matrix MCC = 0.37. (B) Full dataset, right: cut-off and left: confusion matrix MCC = 0.51.

Supplementary Figure 2

FILTERED DATASET

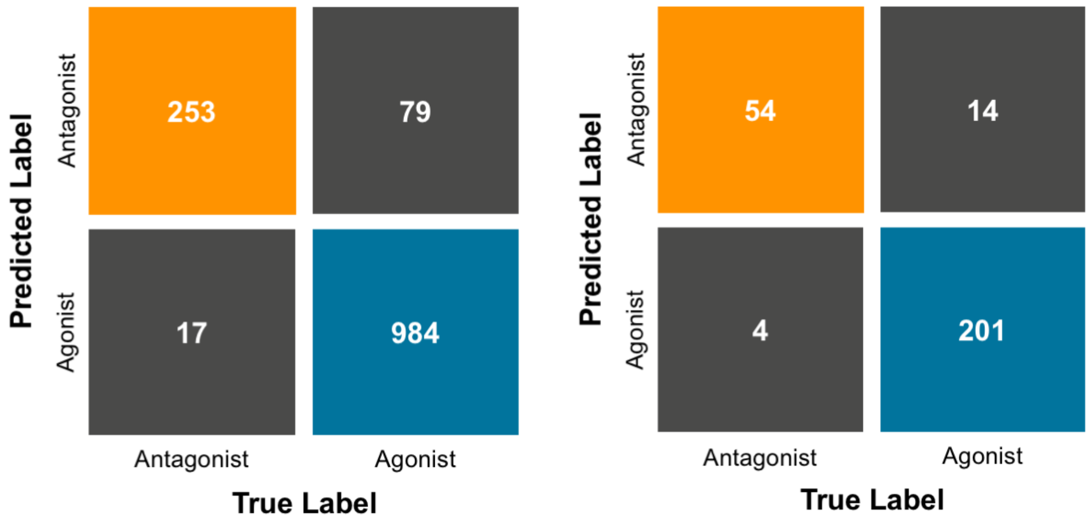


Supplementary Figure 2. Random Forest Classification Confusion Matrix for Cross-Validation (left) and Hold-out sets (right) for the filtered dataset. The left figure shows the median confusion matrix across all K-folds and repeats, the right shows the hold-out set. Y-axis shows the frequency of classes predicted by the model, across all samples. The x-axis shows the true known class label. Overall the confusion matrix provides insight into the ratios of false positive & negatives made by the model across samples for the filtered dataset.

Supplementary Figure 3

FULL DATASET

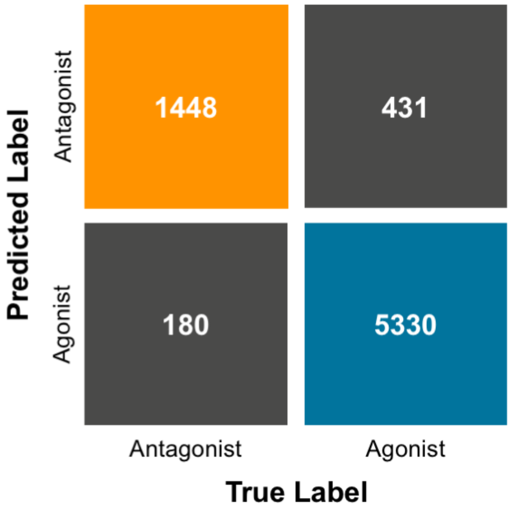


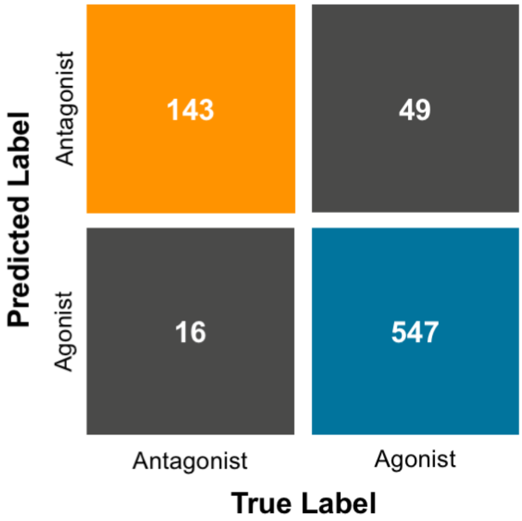


Supplementary Figure 3. XGBoost Confusion Matrix for Cross-Validation (left) and Hold-out sets (right) for the full dataset.

Supplementary Figure 4

A) FILTERED DATASET

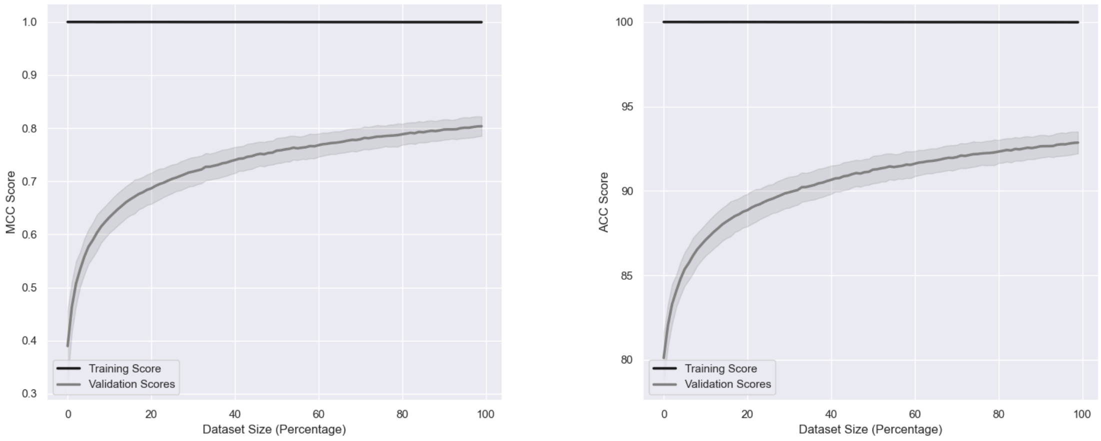


B) FULL DATASET

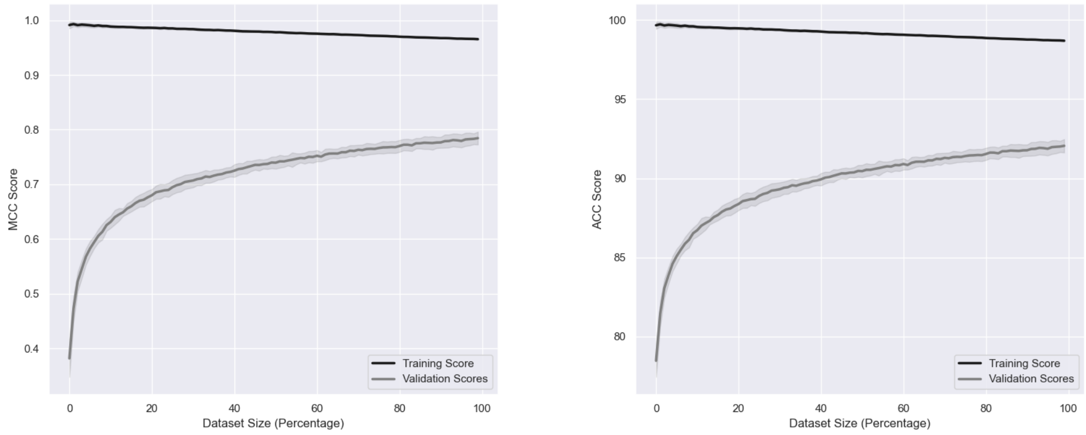


Supplementary Figure 4. Learning curves for the MCC Scores (left) and accuracy – ACC Scores (rights) for the (A) RFC model on the filtered dataset and the (B) XGBoost model on the unfiltered dataset.

Supplementary Figure 5

A

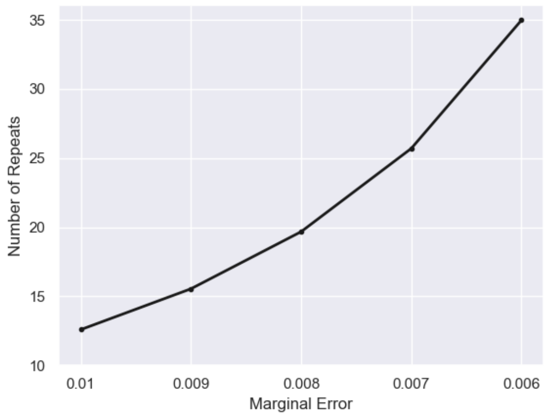


B

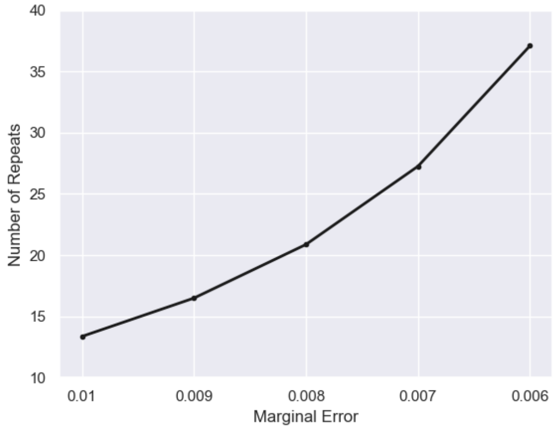


Supplementary Figure 5. Determining the optimal number of repeats required to estimate, with 99% confidence, true model performance to a specified level of precision. (A) RFC model on the filtered dataset and (B) XGBoost model on the unfiltered dataset. In this case, 13 repeats were sufficient for both models.

#### Additional files:

Supplementary Tables 1-5 are in included as an Excel file "SupplementaryTables_1-5.xlsx".
